# Root trait responses to drought depend on plant functional group

**DOI:** 10.1101/801951

**Authors:** Y.M. Lozano, C.A. Aguilar-Trigueros, I.C. Flaig, M.C. Rillig

**Affiliations:** Freie Universität Berlin, Institute of Biology, plant ecology. D-14195 Berlin, Germany; Freie Universität Berlin, Institute of Biology, Applied Zoology/ Animal Ecology D-12163 Berlin, Germany; Berlin-Brandenburg Institute of Advanced Biodiversity Research (BBIB), D-14195 Berlin, Germany

**Keywords:** Grassland ecosystem, global change, plant traits, root diameter, specific root length, specific root surface area, root tissue density, winRhizo

## Abstract

1. Drought can strongly modify plant diversity and ecosystem processes. As droughts are expected to intensify in the future, it is important to better understand plant responses to drought. We expect that roots traits constitute an overlooked but powerful predictor of plant responses as roots are in direct contact with the soil environment, taking up nutrients and water.
2. Here, we determine which root traits are sensitive to drought, the magnitude of that response, whether their predictive power and relationships with shoot biomass are affected by drought and whether all these responses depend on plant functional group. To do so, we conducted a glasshouse experiment with 24 plant species grown in pots (10 replicates per species), which represent three different functional groups: grasses, herbs and legumes. All replicates were well watered during the first month and then half of the replicates were kept under drought (30 % water holding capacity (WHC)) with the other half serving as control (kept at 70% WHC). After two months of the treatment, leaf and root traits were measured.
3. Leaf traits had a strong but more uniform response to drought compared to root traits. Root trait response was variable and differed by plant functional group. Most grasses had increased root diameter, specific root surface area (SRSA) while decreased root tissue density (RTD) with drought. Production of thicker roots with a low tissue density could allow grasses to achieve greater nutrient and water acquisition through mycotrophy and would be linked to an increase in the reserve of non-structural carbohydrates needed for osmoregulation. Herbs had decreased SRSA and specific root length (SRL) while increase root carbon allocation. Reduction of root elongation or sacrifice of fine roots would be compensated by an increase in root carbon allocation, which allow herbs to improve water uptake. Legumes did not alter root morphological traits but promote an early flowering in order to scape drought.
4. Our results identify changes in root morphological traits as mechanisms to likely face drought, a response that is species-specific and differed among functional groups.

## INTRODUCTION

Plant traits provide a means of explaining the role of plant diversity in ecosystem functioning. Given this explanatory power, there has been a lot of interest in measuring the response of plant traits to climatic factors. This research, though, has been biased towards leaf traits, neglecting studies focused on root traits. This bias is unfortunate because root traits are likely better predictors of ecosystem functioning than leaf traits as roots are in direct contact with the soil environment, taking up water and nutrients, and they modify the physical, chemical and microbial properties of the soil (Bardgett, Mommer & De Vries 2014). Indeed, evidence points to root traits driving ecosystem processes such as carbon and nutrient cycling, soil formation and stability (Bardgett, Mommer & De Vries 2014). This lack of research on root traits hampers our ability to predict how plant diversity and ecosystem functioning will respond to climate change. In particular, climate models predict that drought episodes may intensify in the future due to warmer temperatures and potential decline in seasonal rainfall (Bodner & Robles 2017).

Data available on root trait responses to drought is limited to a small number of plant species. As a result, conclusions about plant strategies in term of root trait responses seem idiosyncratic or at best premature. For example, some studies report plant species producing thinner roots with high specific root length (SRL) and specific root surface area (SRSA) in response to drought, a strategy interpreted to improve soil moisture acquisition with a low plant investment (Debinski *et al.* 2010; Comas *et al.* 2013). Other studies, by contrast, report that plant species produce thicker roots with a low SRL and SRSA, which has been shown to diminish the risk of hydraulic rupture (Zimmermann 1983; Zufferey *et al.* 2011). Thicker roots have been associated with high nutrient and water acquisition through mycotrophy (Brundrett 2002; Comas *et al.* 2012), and with osmoregulation due to the storage of non-structural carbohydrates (NSC) (Chaves 1991; Galvez, Landhäusser & Tyree 2011; Yang *et al.* 2016). Recent meta-analyses found that drought decreases root length, and increase root diameter and root:shoot ratio (Zhou *et al.* 2018). However, we argue that the contrasting patterns observed in root trait responses to drought could be better understood when taking into consideration the fact that plant strategies are shaped by their evolutionary history (Reich 2014). Thus, root trait response to drought are likely not generalizable across all plant species. Instead, such responses could potentially be predicted based on plant membership to functional groups, which reflect phylogenetic relationships. Plant functional groups have been adopted by modelers to represent broad groupings of plant species that share similar characteristics and roles in ecosystem function (Wullschleger *et al.* 2014). In fact, the use of plant functional groups has proven to be an efficient way to model how drought modifies plant communities and how that change feedback on the local environment both in current and future climates (Wilson *et al.* 2018). Fast growing species, such as many grasses, which tend to exhibit traits linked with an acquisitive strategy (high SRL, SRSA and low diameter) (Ravenek *et al.* 2016), would modify root traits in a way to overcome drought, although it has been found that some herbs plants also exhibit a similar strategy (Comas *et al.* 2013).

The mechanisms plants have evolved to cope with drought can be categorized into dehydration escape, dehydration avoidance, dehydration tolerance, dormancy, and desiccation tolerance (Volaire 2018). Dehydration avoidance relies on mechanisms that maintain the plant’s water status by decreasing water loss through reduced stomatal conductance and restrict shoot growth, or by maintaining water uptake through physiological, biotic or root morphological adjustments (Brunner *et al.* 2015). Dehydration tolerance is often ascribed to cellular and tissue factors including osmoregulation, and mainly involves increased osmoprotectant levels and accumulation of carbohydrates in plant tissues (Blum 2005; Brunner *et al.* 2015). Although both strategies depend on a combination of different functional traits, dehydration avoidance appears as one of the first strategies plants use to face drought (Volaire 2018); this is when potential shifts in root traits as SRL, SRSA, root tissue density (RTD), root diameter (RAD) or root:shoot ratio become relevant, as roots are the first organ in contact with the soil and thus the first that face water scarcity. In addition to these root morphological traits, drought can also affect leaf and root N content (He & Dijkstra 2014; de Vries, Brown & Stevens 2016). A reduction in soil moisture may reduce microbial activity, net N mineralization (Borken & Matzner 2009), nutrient diffusivity and mass flow (Rouphael *et al.* 2012), which in the end would reduce plant N uptake.

Plant economics spectrum (PES) theory predicts that the leaf, stem and root traits related to resource acquisition are strongly correlated with each other, and this has been proposed to explain ecological strategies for acquiring, processing and retaining multiple limiting resources (Wright *et al.* 2004; Reich 2014; Kramer-Walter *et al.* 2016). From the root economic spectrum (RES) perspective, traits positively associated with water and nutrient uptake capacity, like high SRL, should correlate negatively with root tissue density (RTD) and diameter (Reich 2014). However, evidence for these trends in roots is weak at best and inconsistences with the expected RES trends are common (Valverde-Barrantes & Blackwood 2016). Evidence suggests that root traits, such as SRL and RTD, are independent of each other and from the leaf economic spectrum (Kramer-Walter *et al.* 2016). It seems that the PES may depend on the plant functional group, as for instance, graminoid plants are aligned with the RES syndromes (Roumet *et al.* 2016).

In this study, we measured leaf and root trait responses to drought of 24 plant species belonging to three plant functional groups: grasses, herbs and legumes. With this information, we aimed to determine root trait response to drought, if these are linked to the PES hypotheses, if leaf and root traits responses are coupled, if their predictive power and their relationships with shoot biomass are affected by drought and if all these responses depend on plant functional group.

## MATERIALS AND METHODS

### Species selection

We selected 24 plant species belonging to three different plant functional groups: eight grasses (*Arrhenatherum elatius, Festuca brevipila, Holcus lanatus, Poa angustifolia, Anthoxanthum odoratum, Lolium perenne, Festuca rubra, Dactylis glomerata)*, thirteen herbs *(Achillea millefolium, Armeria maritima ssp.elongata, Artemisia ssp. Campestris, Berteroa incana, Daucus carota, Galium verum, Hieracium pilosella, Hypericum perforatum, Plantago lanceolata, Potentilla argentea, Ranunculus acris, Rumex thyrsiflorus, Silene vulgaris)* and three legumes *(Trifolium repens, Vicia cracca, Medicago lupulina).* All these species are common, frequently co-occurring grassland species in Central Europe. Plant species will be referred to by their generic name from here on (except for the two *Festuca* species to which we refer as *F. brevipila* and *F. rubra*). Seeds of these plant species were obtained from commercial suppliers in the region (Rieger Hoffmann GmbH, Germany).

### Experimental design

In September 2016, we collected sandy loam soil (% N 0.07, % C 0.77, pH 6.66) from Dedelow, Brandenburg, Germany (53° 37’ N, 13° 77’ W) where our plant species naturally grow. Soil was sieved (4 mm mesh size) and homogenized to use as substrate. We established the experiment in a controlled glasshouse growth chamber with a daylight period set at 12 H, 50 klx, and a temperature regime at 22/18 °C day/night with relative humidity of ∼40 %. Prior to germination, we surface-sterilized seeds with 10% sodium hypochlorite for 5 min and 75% ethanol for 2 min and then thoroughly rinsed with sterile water. Then, seeds were germinated in trays with sterile sand and transplanted into deep pots (11 cm diameter, 30 cm height) five days after germination. Pots were filled with 3 L of soil and one individual seedling per plant species was planted into the center of each pot (for a total of 10 replicate pots per plant species). Thus, this experimental design included twenty-four plant species x two water treatments x five replicates = 240 pots. Plants that died during the first two weeks were replaced.

All plants were well-watered during the first month of growth. Then, half of the pots (i.e. 5 replicates of each plant species) were kept under drought treatment by maintaining 30% water holding capacity (WHC) while the other half were maintained as control at 70% WHC for two months. Pots were weighed every two days and their moisture content adjusted gravimetrically with an accuracy of ± 0.5 g to keep them at their respective WHC during two months. All pots were randomly distributed in the chamber and their position shifted twice to homogenize environmental conditions during the experiment.

At harvest, we clipped the plant aboveground material while the belowground compartment, consisting of both soil and roots, of each pot was divided into three sections every 10 cm of depth (upper, middle and bottom sections). The soil and roots from these sections were pre-dried at 27 °C in an incubator for three weeks to stabilize samples. After that, we first collected roots by hand. Then, we spread the soil on a filter paper to capture small and fine roots (less than 1 mm) by electrostatic. This was done by manually collecting root pieces attracted to an electrostatically charged polyethylene plate (we charged the plate by means of intensive rubbing for 4-5 seconds on a stretched wool fabric) in order to ensure the capture of all fine roots. We used this capture method on dried root sample for two reasons. First, this method allows for better recovery of fine root pieces (as opposed to root washing techniques where small and fine roots are easily lost). Second, it allows fast storage of root sample until further processing (see below). This latter point was important because given the large amount root samples it was not feasible to measure root traits on fresh samples without risking root degradation.

### Measurements

#### Plant morphological traits

We measured the traits in fine roots (i.e., < 2 mm in diameter): length, surface area, volume and root average diameter on a rehydrated root sample from the middle section of the belowground compartment (i.e. the section in between 10-20 cm depth) using the WinRhizo™scanner-based system (v.2007; Regent Instruments Inc., Quebec, Canada). We focused our analysis on the middle section of the soil because we did not find differences in root traits due to root section (e.g., *Anthoxanthum, Daucus*). This method provides accurate and unbiased root trait values as shown in previous studies that report high linear correlations (Pearson’s r = 0.93) between root traits measured on fresh and rehydrated material of these species (Bergmann *et al.* 2017).

#### Biomass traits

We measured root and shoot mass after drying samples at 70 °C for 48 h.

#### Aboveground traits

We also measured specific leaf area (SLA) and leaf dry matter content (LDMC) following standard protocols (Cornelissen *et al.* 2003).

#### Chemical and physical traits

We measured soil, leaf and root C and N contents with an Elemental Analyzer (EuroEA, HekaTech, Germany). Soil temperature (Hobo 1-800, Onset Computers, Pocasset,MA, USA) at depth of 15 cm were monitored continuously during the experiment in additional control pots under drought and non-drought conditions.

### Statistical analyses

#### Plant trait responses to drought

First, we tested whether the drought treatment affected all plant traits (aboveground and belowground) depending on plant species identity using two-way MANOVA (i.e. drought treatment and plant species identity were used as main factors). Then, we used linear models to determine which plant traits significantly responded to drought treatment and whether such responses were consistent across plant species. We use a two-way ANOVA that reflected our full factorial design with water availability (drought and non-drought), plant species (i.e., 24 species) and their interactions, considered as fixed factors on each plant trait separately. We include in our models the correlation among species within each functional group by using the correlation structure function (corSymm) and soil temperature as covariate. Leaf (shoot mass, SLA, LDMC, leaf C and N) and root traits (root average diameter (RAD; mm), root tissue density (RTD; root dry weight per volume mg cm^-3^), specific root length (SRL; cm mg^-1^), specific root surface area (SRSA; cm^2^ mg^-1^), root to shoot ratio, total root mass, root C and N) were transformed to meet homoscedasticity and normality assumptions. Analyses were conducted using R version 3.5.3 (R Core Team 2019). Additional analyses were done using the R interface implemented in Infostat-Statistical Software (Di Rienzo *et al.* 2017). Results shown throughout the text and figures are mean values ± 1 SE.

Second, we determined the direction and magnitude of the effect of drought on each plant trait by using the relative interaction index (RII, (Armas, Ordiales & Pugnaire 2004). This index provides a means to determine whether trait values are higher or lower under drought relative to non-drought (control) conditions. It is calculated as ratio as follows:

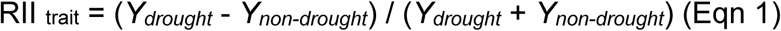

Where *Y*_*drought*_ is the response of the trait when the plant grew under drought (30% WHC) and *Y*_*non-drought*_ is the response of the trait when the plant grew under non-drought conditions (70% WHC). The index was calculated for each plant species and for each plant functional group. RII has positive values when trait values are greater under drought than in non-drought and negative when the opposite is true. In addition, we performed a priori contrasts tests to assess whether RII values were significantly different from zero (indicating neutral or no significant effect).

Finally, we determined whether there was a higher variability in the plant trait response under drought or under non-drought conditions by evaluating the standard deviation of the trait. We also identify the root traits that better predict shoot mass and analyze the relationships between root traits and shoot mass under each water condition. Graphical inspection via radar charts, Pearson’s correlations, regressions and principal components analyses were performed.

## RESULTS

### Plant species differ in the expression of leaf and root traits depending on the plant functional group

In terms of leaf traits, ordination by PCA showed a clear separation between grasses and herbs/legumes (Fig 1a), with the first two axes explaining 61.7 % of the total variance under non-drought conditions. Contrary to expectations, grasses, also known as “fast growing species” showed a higher shoot mass, leaf dry matter content but a lower SLA and leaf N than the other two plant functional groups. In terms of root traits, PCA showed a clear and better separation between plant functional groups compared to leaf traits (Fig. 1b). The first two axes explained 70.4 of the total variance under non-drought conditions. Grasses showed greater SRL and SRSA but a smaller root diameter and RTD in comparison with herbs and legumes. Legumes showed a higher root N content and a lower root mass production than grasses and herbs

**Figure 1.**
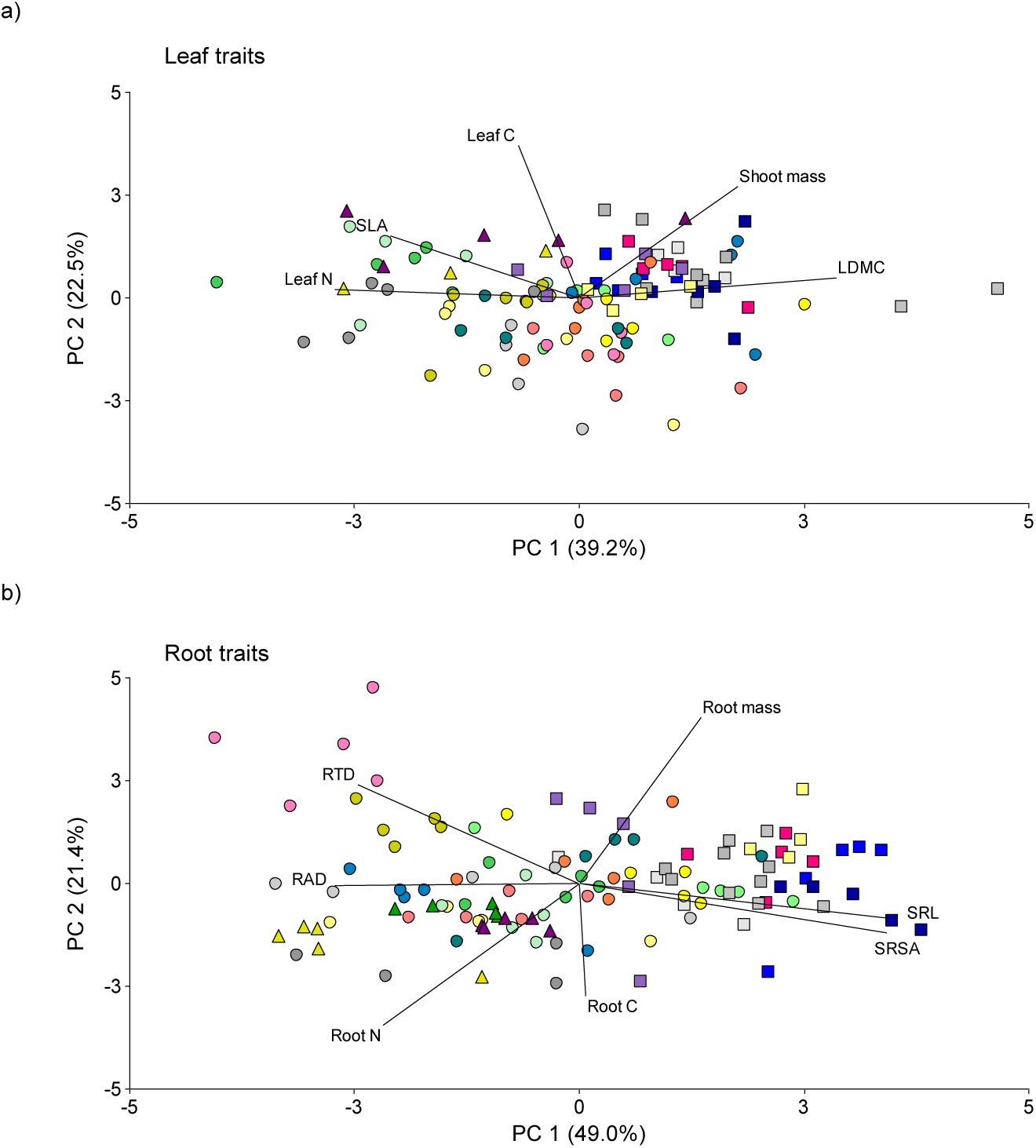
Principal component analysis of leaf (a) and root (b) traits associated to 24 plant species belonging to different plant functional groups growing under non-drought conditions. Plant functional groups: Grasses (square symbols), herbs (circle symbols) and legumes (triangle symbols). Plant traits: SLA (specific leaf area), LDMC (leaf dry matter content), RDA (root diameter), RTD (root tissue density), SRL (specific root length), SRSA (specific root surface area). Plant species: *Anthoxanthum* (blue 2), *Berteroa* (light yellow), *Daucus* (light green), *F.brevipila* (light gray), *Achillea* (salmon), *Potentilla* (deepskyblue4), Plantago (yellow), *Hieracium* (darkseagreen1), *Artemisa* (azure 2), *Holcus* (pink), *Arrhenaterum* (purple), *Vicia* (golden), *Hypericum* (chartreuse), *F.rubra* (azure 3), *Galium* (orange), *Poa* (dark blue), *Silene* (dark golden), *Trifolium* (green), it was not included in the graph ‘a’, *Dactylis* (azure 4), *Rumex* (light pink), *Medicago* (dark purple), *Ranunculus* (Cyan 4), *Armeria* (Dark grey). n=5.

### Root trait responses to drought are more heterogeneous than leaf trait responses

Although both leaf and root traits values were affected by drought, species identity and their interaction (Table 1, S1, S2), the relative interaction index (RII) of leaf traits was homogenously negative across plant species (Fig. 2). That is, shoot mass, SLA, and leaf N all had lower values under drought relative to the non-drought treatment while LDMC had an increase in response to drought (Fig. 2a, b, d, e). In contrast, most root traits were more heterogenous in their response to drought, depending on plant species identity. For example, grasses such as *Dactylis* and *Arrhenaterum* and the herbs *Hypericum, Daucus* or *Achillea* produced more root biomass under drought relative to non-drought, while we observed the opposite pattern for the legumes *Vicia, Trifolium* and *Medicago*, the grasses *Poa, Lolium, F.rubra, F.brevipila* and the herbs *Rumex, Ranunculus* and *Galium* (Fig. 2f). Root diameter was greater under drought compared to non-drought in most species except *Trifolium, Silene* and *F.brevipila* (Fig. 2g) while the opposite was true for RTD (Fig. 2i). Root-shoot ratio increased under drought for most herbs and for some grasses as *Dactylis* and *Arrhenaterum* (Fig. 2h). SRL was lower under drought compared to non-drought in most herbs, except *Hieracium* or *Silene* (Fig. 2j), while for grasses and legumes, some species had the opposite pattern. SRSA was lower in drought compared to non-drought for most herbs (i.e., *Rumex, Galium, Artemisia, Armeria*), while the opposite was true for most grasses (i.e., *F.rubra, F.brevipila, Arrhenaterum, Anthoxanthum*). The legumes *Vicia* or *Medicago* had lower SRSA under drought while *Trifolium* had the opposite pattern (Fig. 2k). Root C was higher under drought for most herbs, but a contrary pattern was found in *Poa* (Fig. 2l). Root N was higher under drought for some grasses, and some herbs as *Rumex* or *Daucus* but it was lower for most herbs species as *Silene, Ranunculus, Hieracium* or *Achillea* (Fig. 2m).

**Table 1.** Results from general linear models on leaf and root traits response to water availability (i.e., drought). Leaf dry matter content (LDMC), specific leaf area (SLA), root average diameter (RDA), Root tissue density (RTD), specific root length (SRL), specific root surface area (SRSA). Plant species (Ps), water availability (Wa) and their interactions were considered as fixed factors (table above). F-values and significance (*, p < 0.05; **, p < 0.01; ***, p < 0.001; all significant values are shown in bold; n.s: non-significant).

| Source of variation | Leaf traits |  |  |  |  | Root traits |  |  |  |  |  |  |
| --- | --- | --- | --- | --- | --- | --- | --- | --- | --- | --- | --- | --- |
|  | Shoot mass | LDMC | SLA | Leaf C | Leaf N | Root mass | RAD | RTD | SRL | SRSA | Root N | Root C |
| Plant species (Ps) | **24.15\*\*\*** | **36.81\*\*\*** | **26.34\*\*\*** | **9.65\*\*\*** | **86.67\*\*\*** | **53.07\*\*\*** | **29.42\*\*\*** | **39.42\*\*\*** | **136.23\*\*\*** | **147.51\*\*\*** | **101.34\*\*\*** | **5.63\*\*\*** |
| Water availability (Wa) | **48.43\*\*\*** | **12.08\*\*\*** | **10.63\*\*\*** | 0.20 n.s | **15.66\*\*\*** | 3.11 n.s | **8.95\*\*\*** | **10.42\*\*\*** | 0.68 n.s | 0.46 n.s | 0.19 ns | 1.49 n.s |
| Ps x Wa | **1.97\*\*\*** | **2.29\*\*\*** | **1.71\*** | **1.57\*** | **2.87\*\*\*** | **2.52\*\*\*** | **2.79\*\*\*** | **4.08\*\*\*** | **1.81\*\*** | **2.32\*\*\*** | **3.43\*\*\*** | **2.24\*\*\*** |

**Figure 2.**
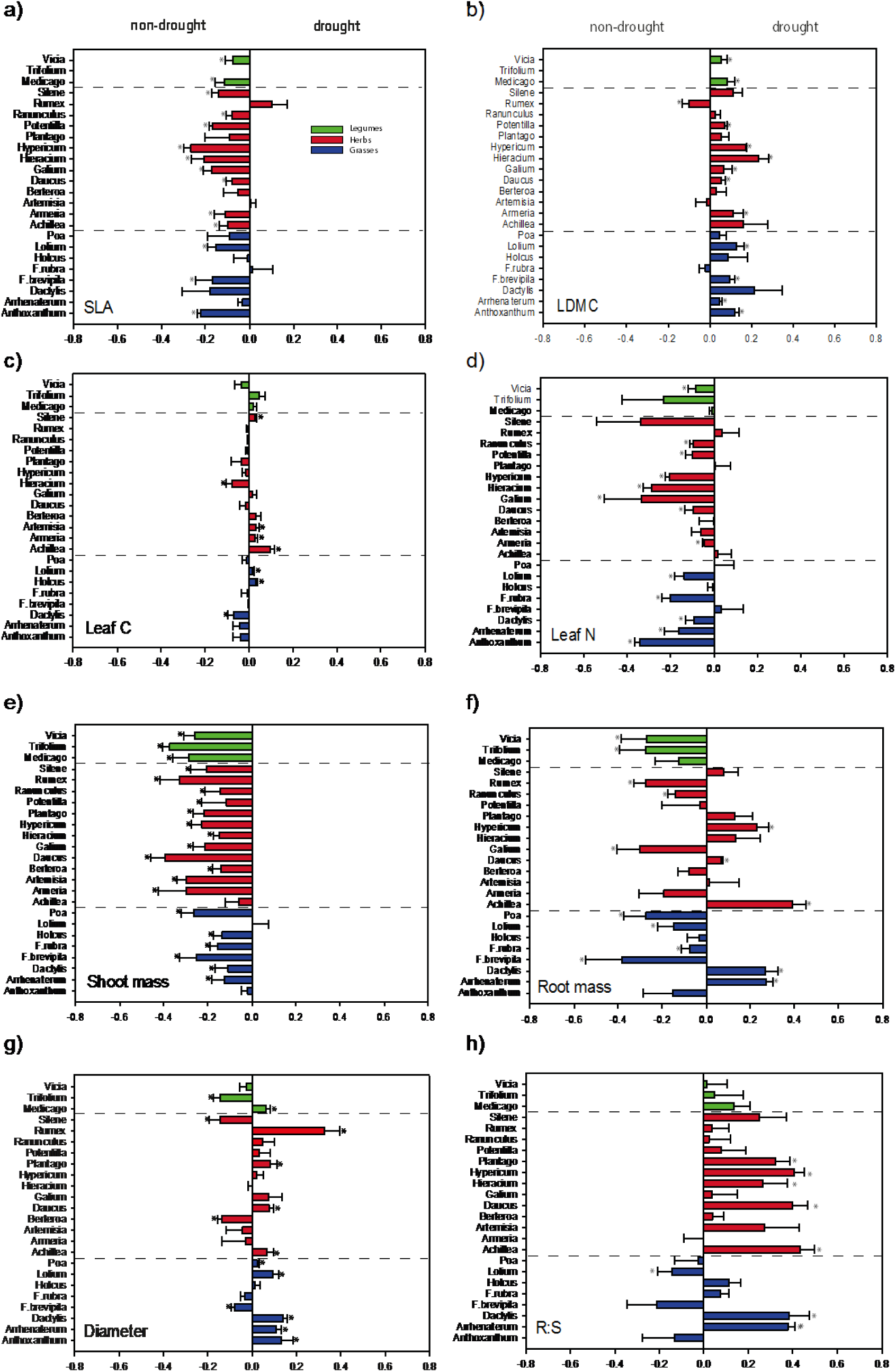
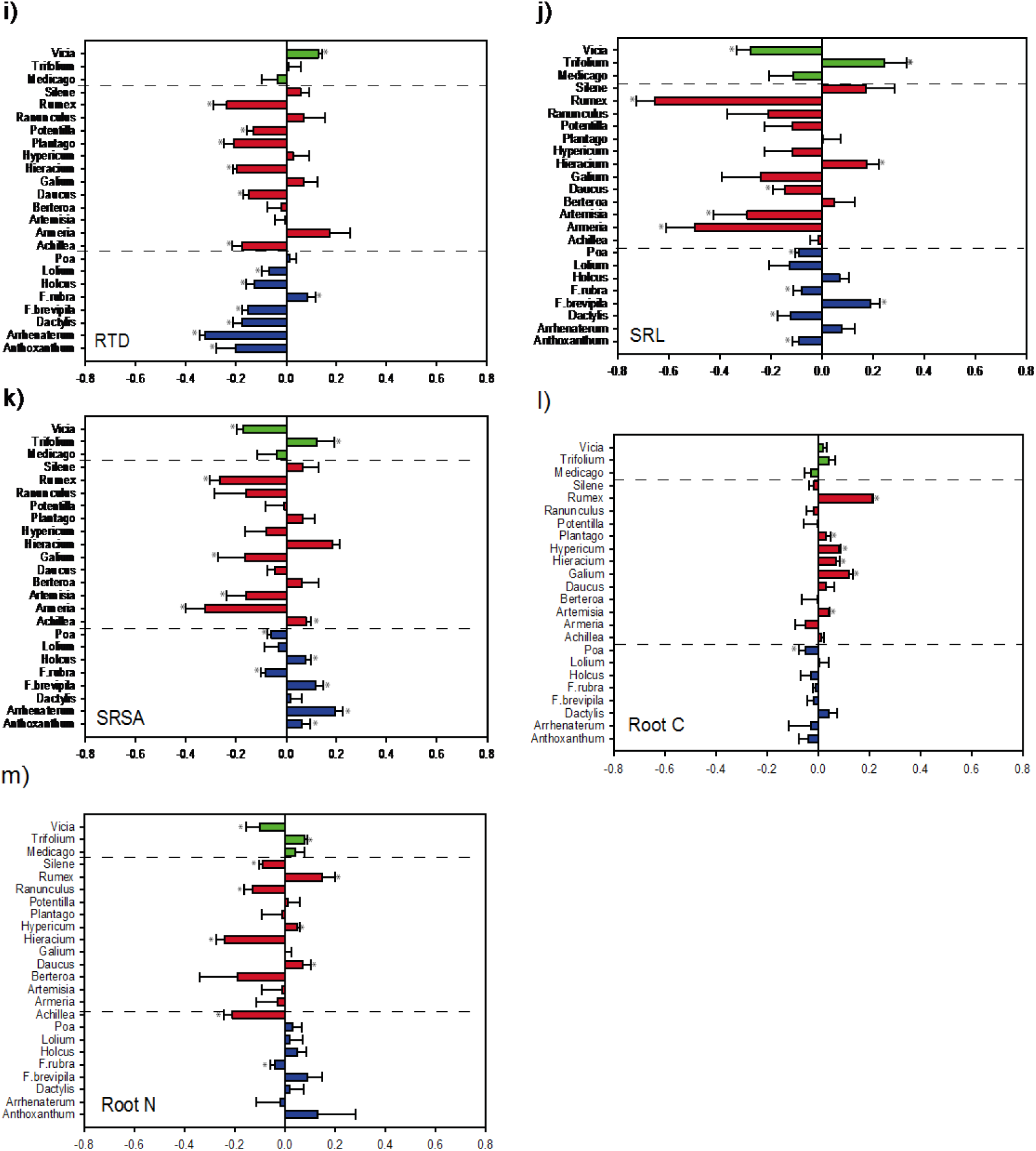
Magnitude of drought effect on leaf and root traits for each plant species. RII compares the trait value in the drought vs non-drought situation. Negative values indicate higher trait value in non-drought than in drought treatment and positive values indicate the opposite. RII values different significantly from 0 are indicated by asterisk (*, P < 0.05).

### Root trait response to drought depended on plant functional group

Root trait response to drought was influenced by plant functional group (Table S3, Fig. 3). Root diameter was greater under drought relative to non-drought for most species, but it was highly evident for grasses and herbs (Fig.3a). Root mass was lower under drought, especially for legumes (Fig. 3b); while SRSA and SRL were lower under drought for most herbs (Fig. 3 c, d). Interestingly, contrary to herbs, higher SRSA under drought was common for most grasses (Fig. 3c). Root tissue density was low for herbs and grasses (Fig. 3e), while root:shoot ratio increased with drought for herbs (Fig. 3f) A summary of the leaf and root trait response is shown in Figure 4.

**Figure 3.**
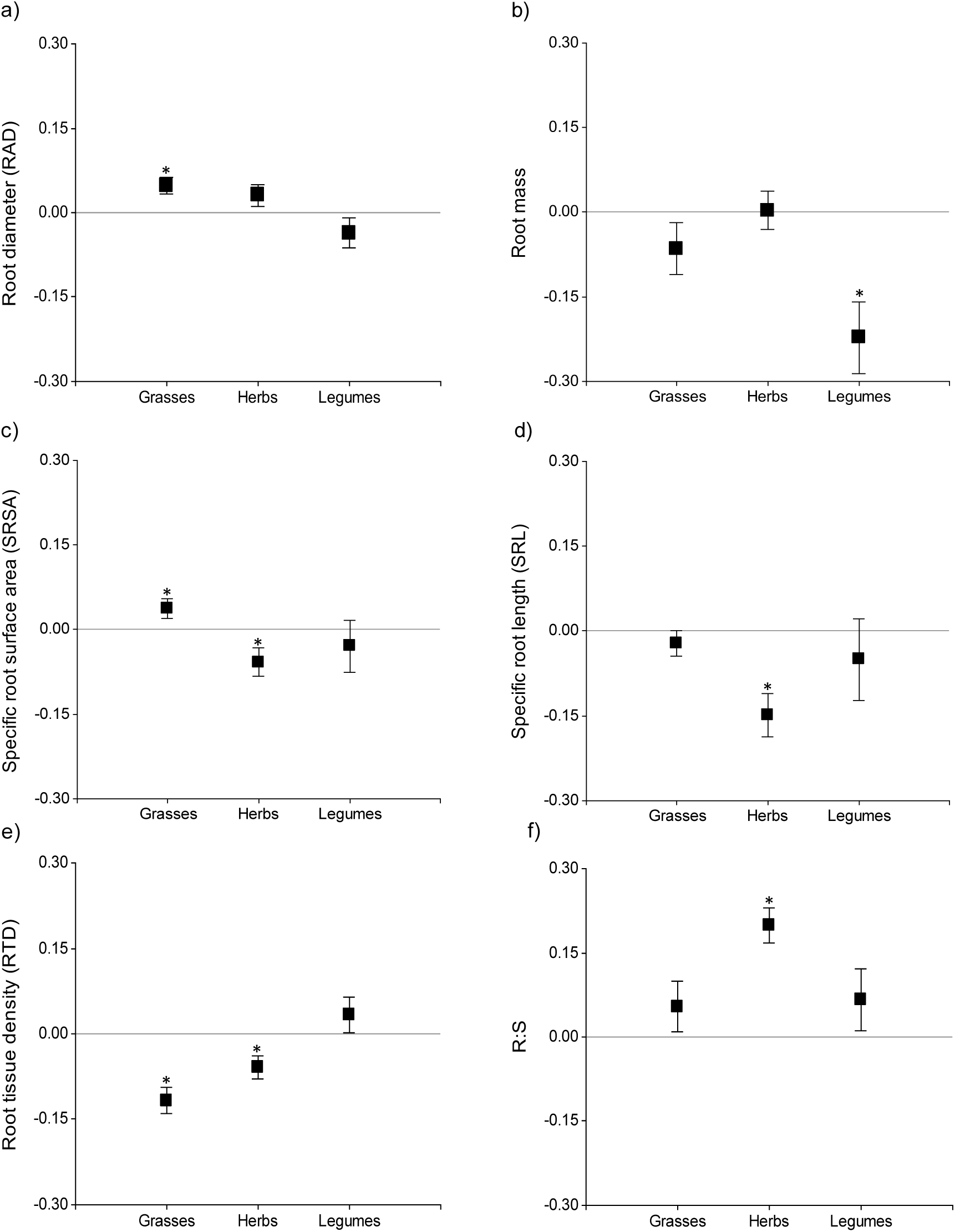
Magnitude of the drought effect on root traits for each plant functional group. Relative index of interaction (RII) compares the magnitude of the effect of drought vs non-drought treatment. Negative values indicate higher trait response in non-drought than in drought treatment and positive values indicate the opposite. RII values different significantly from 0 are indicated with asterisks *, p < 0.05).

**Figure 4.**
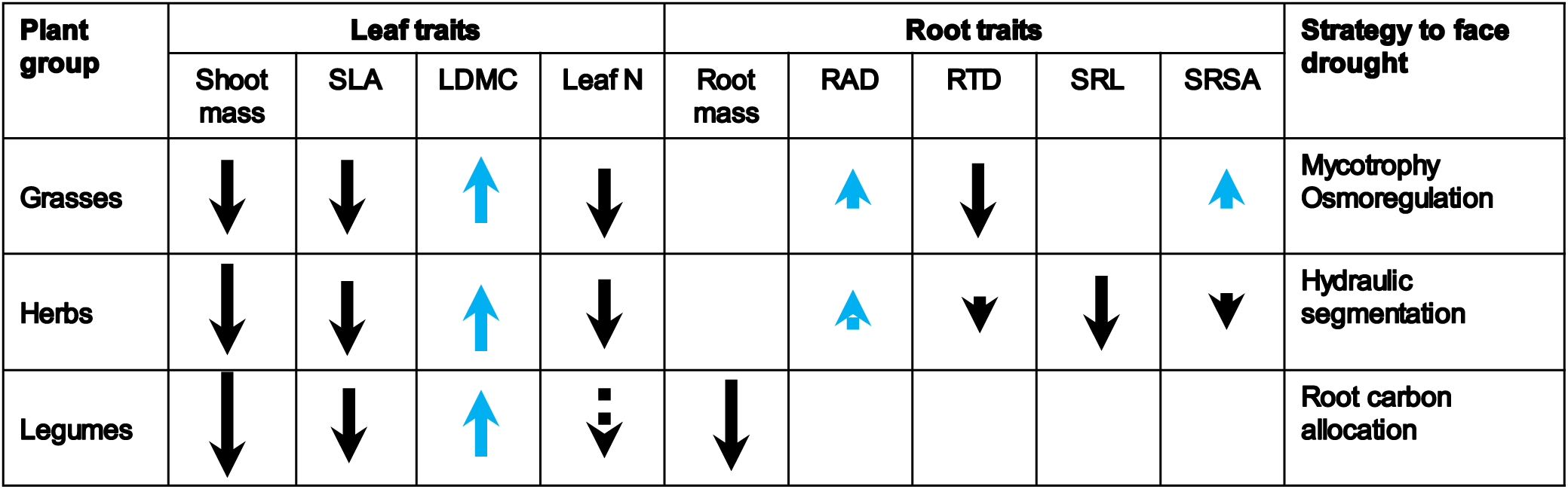
Summary of the plant trait responses to drought. Specific leaf area (SLA), leaf dry matter content (LDMC), root diameter (RAD), root tissue density (RTD), specific root length (SRL), specific root surface area (SRSA). An arrow up indicate higher values in drought than in non-drought treatments (blue), and arrow down indicates the opposite (black). Size of the arrow indicates the magnitude of the effect. Dotted arrow indicates differences at p=0.1, solid arrows at p<0.05. No arrow indicates no differences among drought and non-drought treatments.

### Root trait-shoot biomass relationships were stronger under drought than under non-drought conditions: correlation of traits to shoot biomass and trait variability

Under non-drought conditions, different plant traits were associated with shoot mass but under drought, root traits were best associated and the five species with greatest growth under drought had similar root trait responses (Fig. 5). For example, the five plant species with a higher shoot mass under non-drought conditions (i.e., the herbs *Armeria, Artemisia, Daucus, Rumex* and the legume *Trifolium*), showed no consistent patterns across their root traits (Fig. 5a). *Rumex* showed a high RTD and root mass and a low SRL and root C; *Daucus* showed a high SRSA and SRL values, while *Armeria* a high RAD and root N values. By contrast, under drought conditions (Fig. 5b) the five plant species with higher shoot mass (i.e., the grasses *Anthoxanthum, Dactylis, Lolium* and the herbs *Achillea* and *Potentilla*) showed a low variation in the seven root traits (except *Anthoxanthum* that had a higher SRSA and SRL in comparison to the other plant species). Overall, when considering the 24 species, we observed a lower variability under drought than under non-drought for most of the traits (Fig. S1). In addition, we found a greater correlation between root traits and shoot biomass under drought, than under non-drought conditions (Fig. S2). For instance, R^2^ increased from 0.05 in non-drought to 0.12 in drought, for RAD, (Fig. S2a). A similar pattern was found for RTD, SRL and SRSA (Fig. S2 b-d). Pearson correlation between shoot mass and root traits was higher under drought than under non-drought conditions (Table S4). Root traits that better predicted shoot mass were root diameter, SRL and SRSA, especially under drought conditions (Table 2).

**Table 2.**
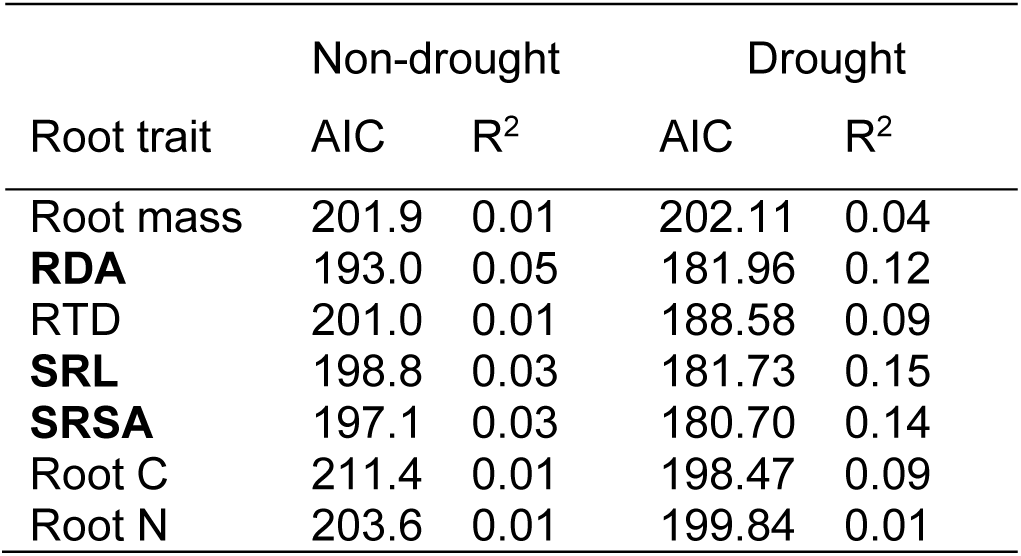
Selection of the best predictors of shoot mass under non-drought and drought conditions. Properties of each treatment were tested in separate models. The fit of univariate linear models is shown with Akaike Information Criterion (AIC) and coefficient of determination (R^2^). Best predictors of shoot mass are marked in bold.

| Root trait | Non-drought |  | Drought |  |
| --- | --- | --- | --- | --- |
| | AIC | $R^2$ | AIC | $R^2$ |
| Root mass | 201.9 | 0.01 | 202.11 | 0.04 |
| **RDA** | 193.0 | 0.05 | 181.96 | 0.12 |
| RTD | 201.0 | 0.01 | 188.58 | 0.09 |
| **SRL** | 198.8 | 0.03 | 181.73 | 0.15 |
| **SRSA** | 197.1 | 0.03 | 180.70 | 0.14 |
| Root C | 211.4 | 0.01 | 198.47 | 0.09 |
| Root N | 203.6 | 0.01 | 199.84 | 0.01 |

**Figure 5.**
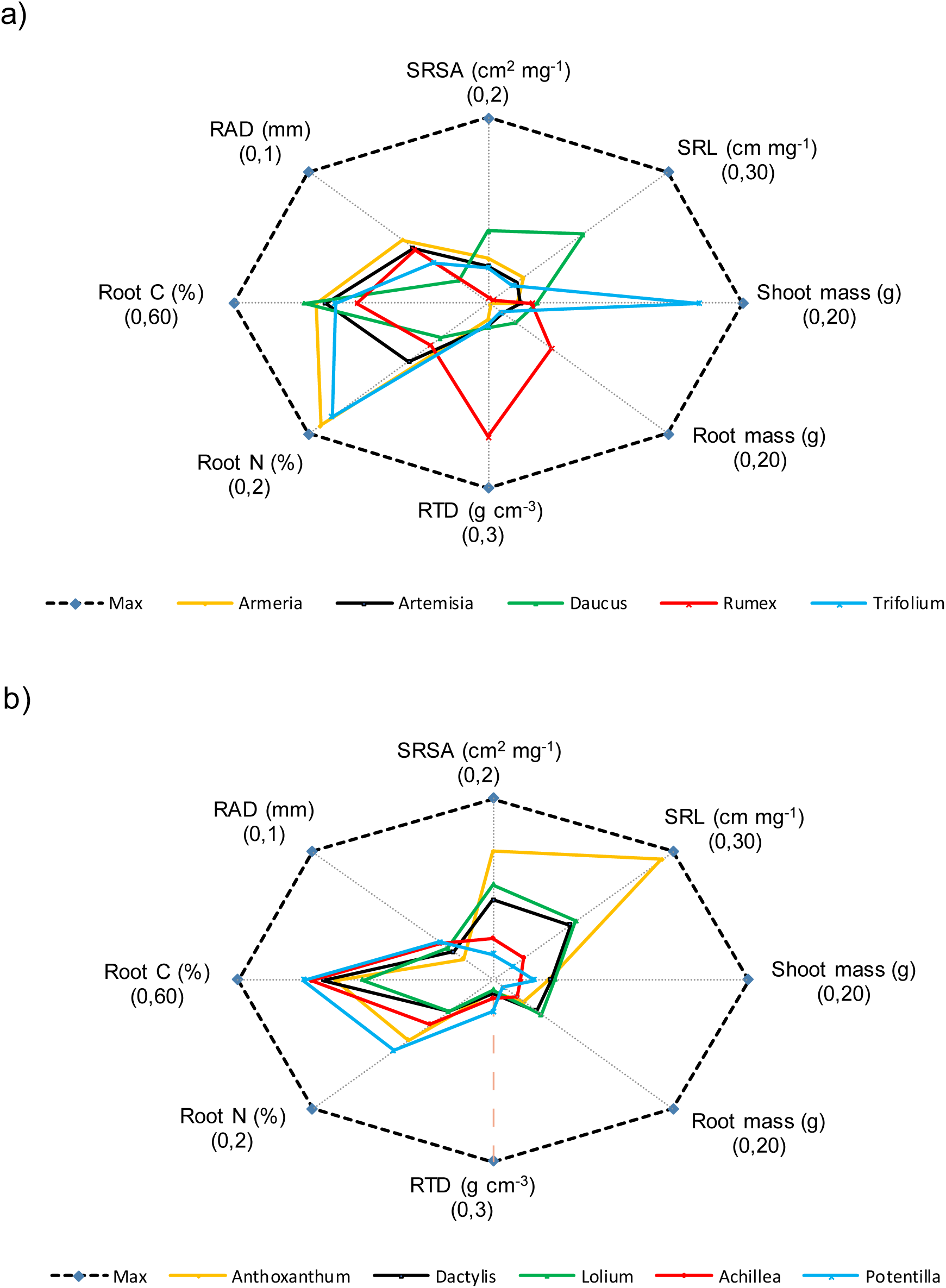
Radar chart of the plant traits of the top five plant species that had the highest shoot mass under a) non-drought and b) drought conditions. Root diameter (RAD), specific root length (SRL), specific root surface area (SRSA), root tissue density (RTD). The scale (0,X) represents the minimum and maximum value for each trait.

In addition to plant traits, soil nutrients were analyzed (Table S5). We found that soil C and N content was only affected by drought for legumes species; they were lower under drought than under non-drought conditions.

## DISCUSSION

The plant economic spectrum has been used to describe general coordinate syndromes (Wright *et al.* 2004; Reich 2014), however, also some inconsistencies have become apparent (Kramer-Walter *et al.* 2016; Valverde-Barrantes & Blackwood 2016). Our results show that PES may depend on the plant functional group. We observed a strong relationship between “fast” traits, such as SLA and leaf N, which are linked with the photosynthetic rate and with a rapid acquisition of key resources (Grime *et al.* 1997), but these ‘fast’ traits were more associated with herbs and legumes than with the “fast growing” grasses species. However, in terms of the RES, ‘fast’ root traits, such as SRL and SRSA, were highly associated with grasses, more so than with herbs or legumes. Under drought conditions, similar patterns than under non-drought were found (Fig. S3).

Leaf and root traits differed in their response to drought. Leaf traits had a stronger response to drought than root traits and it was consistent across plant species (i.e. independent of the plant functional group). For example, all plants species under drought had reduced leaf area (which is correlated with decreased photosynthetic activity) and increased leaf thickness (which would be indicative of more rigid cell walls (Markesteijn *et al.* 2011). These responses enable leaves to maintain turgor and minimize cell damage under drought and has been related to an increase in drought survival (Bongers *et al.* 2017). The decrease in SLA, which is strongly correlated with LDMC, is associated with drought tolerance (Markesteijn *et al.* 2011). The higher impact on drought on leaves than on root traits may be linked with the fact that roots are the organs responsible of taking up water and the first responders to many kinds of stress (Brunner *et al.* 2015; Weemstra *et al.* 2016), so that, due to their ability to grow toward wetter patches in the soil (hydrotropism, (Eapen *et al.* 2005), roots will be first beneficiaries of the scarce water. In addition, the amount of water that reaches the leaves will be diminished as roots require water for their metabolism (Bais *et al.* 2001) and as the pressure in the xylem becomes excessive under drought, which breaks the water column (cavitation) (Zimmermann 1983). However, although plants would develop strategies to avoid cavitation and embolisms (Zufferey *et al.* 2011), the loss of water through transpiration, as they need to open stomata to absorb CO_2,_ is extremely high (Beerling & Franks 2010).

By contrast, root trait responses depended on the plant functional group so that our results showed that leaf and root trait responses to drought are uncoupled. It is still unclear whether plants are more flexible to alter their root morphology or root biomass allocation under stressful conditions. Freschet, Swart and Cornelissen (2015) have shown that root mass turned out to be far more important than morphological traits under stressful conditions (e.g., nutrients and light), but see Poorter and Ryser (2015). Our results evidence that the change in root biomass allocation or root morphological traits as response to drought depends on the plant functional group: most grasses and herbs mainly rely on morphological root trait adjustment, while only herbs evidence a change in root biomass allocation. Legumes did not change root morphological traits or root biomass allocation.

Plants can modify root traits as a passive response to stress conditions or as an active adaptive response to face drought as seen for the plant osmotic adjustment (Sanders & Arndt 2012), but a different experimental setup is needed to disentangle this issue. Our results showed that grasses had increased root diameter and SRSA and reduced RTD as response to drought. Roots with lower tissue density may support faster acquisition but maintain a shorter lifespan (Wahl & Ryser 2000; Withington *et al.* 2006). We found that RTD is negatively correlated with root diameter. This could be because root cortex is less dense than root stele and because in thicker roots a larger proportion of the root cross-sectional area is accounted for by the cortex (Chen *et al.* 2013; Kong *et al.* 2014). On the other hand, thicker roots may acquire resources faster because of their greater dependence on mycorrhizal fungi (Brundrett 2002; Weemstra *et al.* 2016; Kong *et al.* 2017), but see Maherali (2014) who did not find a correlation between thick roots and fungal colonization.

In addition, thick roots are an important reserve of non-structural carbohydrates (NSC) (Guo, Mitchell & Hendricks 2004). NSC are free, low molecular weight sugars that underpin unique adaptive strategies to environmental shocks (Wang *et al.* 2017). NSC can be utilized to maintain a necessary concentration of soluble sugars (osmolytes) needed for osmoregulation and osmoprotection (Chaves 1991). Thus, the apparent disadvantage of building an expensive root system with thick roots under limited water conditions, may be further compensated by living longer (Kong *et al.* 2017), as afforded by osmotic adjustment using NSC. In fact, there is evidence that under drought conditions thick roots increased NSC accumulation (Yang *et al.* 2016). However, whether the increase in root diameter due to drought is directly linked with an accumulation of NSC remains to be tested. In summary, building thick roots with high SRSA appears to be a key strategy for grasses to tolerate drought. Studies that included a couple of grasses (i.e., *Anthoxanthum* and *Dactylis*) which showed a higher diameter with drought (de Vries, Brown & Stevens 2016), agree with the common pattern we found for grasses.

In herbs, on other hand, their low SRSA and SRL values indicate that this functional group reduces the amount of fine roots. In fact, our results showed that length of roots with a diameter lower than 0.1 mm was lower under drought than under non-drought conditions (Fig. S3). This response can be interpreted as a drought coping mechanism when taking into consideration water flow dynamics. On the one hand, as soon as water availability decreases, the turgor potential of the plant cells diminishes; as a consequence, many turgor-driven processes, such as root elongation, slow down (Bardi & Malusà 2012). On the other hand, the sacrifice of fine roots may prevent the propagation of embolism in the plant, which diminishes the risks of hydraulic rupture under drought (Zufferey *et al.* 2011). This process follows the “hypothesis of hydraulic segmentation”, which states that distal organs, in our case fine roots, are more vulnerable to embolisms (Zimmermann 1983). The reduction in fine absorptive roots also could explain the decrease in root and leaf N under drought conditions. In addition, we did not find any difference in soil N between drought and non-drought treatments, contrary to de Vries, Brown and Stevens (2016), who found differences in dissolved organic N for the herb *Rumex.* It seems that herbs were no able to take up this nutrient when soil moisture decreased. This is in agreement with He and Dijkstra (2014), who demonstrates that drought has a stronger negative effect of plant N uptake. For herbs, the response to drought rely on the increase of conservative root traits (i.e., thick diameter, low SRL and SRSA) and on the increase of root carbon allocation. Some studies suggest that the proportional allocation of carbon to roots increases under drought in order to promote water and nutrient acquisition (Palta & Gregory 1997; Burri *et al.* 2014), which can be linked with an increasing requirement for osmotically active C compounds under drought (Chaves, Maroco & Pereira 2003) On the other hand, it has been commonly observed that root respiration declines under drought (Huang & Fu 2000; Thorne & Frank 2009); and Hasibeder *et al.* (2015) found that drought reduced the amount and the speed of carbon allocation to the root biomass by c. 50%. Nevertheless, our results agree with a recent meta-analyses (Zhou *et al.* 2018) which argue that drought decreases root:shoot ratio.

Finally, legumes reduced root biomass but did not alter root morphological traits or root carbon allocation in response to drought. Reduction in root biomass may be because legumes were dying from hydraulic failure or because they prefer to invest more to photosynthetic or reproductive tissues. We observed an early flowering under drought (e.g., *Trifolium* had 2 ± 1.2 flowers under drought *vs* 0 flowers under non-drought, after 1 week with drought) which is a well-known drought escape mechanism in plants (Shavrukov *et al.* 2017). In addition, it has been suggested that a decrease in root biomass would increase soil N because of the reduction in N root uptake (de Vries, Brown & Stevens 2016), but we found the opposite pattern: a decreased root biomass in legumes linked with a decrease in soil N and a trend to increased root N content. Plants can adjust their uptake kinetics to compensate and facilitate N uptake (Bassirirad 2000) or due to symbiotic N-fixing associations typical of legume species. Legumes establish positive association with bacteria and mycorrhiza which could enhance their drought tolerance (Püschel *et al.* 2017; Naylor & Coleman-Derr 2018), as it has been seen that some bacterial strains promote plant growth under drought presumably due to the production of auxins (Wang *et al.* 2014). A higher N uptake will decrease soil N content. However, although the fixed nitrogen is in the plant, the reduction in root biomass would decrease the biomass of N-fixers and thus the amount of N release to the soil when bacteria died, and/or the amount of nitrogen that can be “leaked” or transferred into the soil (∼3706–6177 kg km^-2^) (Walley *et al.* 1996).

Overall, root trait response was less variable under drought than under non-drought conditions. Stressful conditions constrained the response of root traits while well-watered conditions allow a much more variable trait expression. This can be linked with the fact that the species that better performed under drought are naturally present in dry and semi-dry grasslands (i.e, *Dactylis*, (Federal Agency for Nature Conservation 2019) while species that better performed under non-drought conditions are frequently found in a variable range of environments (i.e., *Trifolium*).

We conclude that root trait responses to drought depend on the plant functional group while leaf traits response is consistent across functional groups. Grasses develop thicker, which would allow for a greater nutrient and water acquisition through mycotrophy, and could promote the accumulation of non-structural carbohydrates needed for osmoregulation. Herbs reduce fine roots (low SRL and SRSA) and increase root carbon allocation. Fine roots can decrease due to reduction in root elongation or as a strategy to prevent the propagation of embolisms in the plant. Increase in root allocation will promote water and nutrient uptake. In contrast to grasses and herbs, legumes do not change root morphological traits or root C allocation. It seems that they promote early flowering, a well-known strategy to escape drought. Root diameter, SRL and SRSA appear as the root traits most strongly linked with plant productivity, and are thus likely candidates for further monitoring. Our result support the inclusion of root traits and their role in ecosystem functioning in future models in order to better project responses of terrestrial ecosystems to global change, specifically to drought.

## ACKNOWLEDGMENTS

The work was funded by the German Federal Ministry of Education and Research (BMBF) within the collaborative Project “Bridging in Biodiversity Science (BIBS)” (funding number 01LC1501A). The authors declare that there is no conflict of interest.

## DATA ACCESIBILITY

Data on plant and root traits: uploaded as online supporting information.

## AUTHORS’ CONTRIBUTIONS

YML, CAAT and MCR conceived the ideas and designed methodology; YML, ICF and CAAT established and maintained the experiment in the greenhouse; YML collected, analyzed the data and led the writing of the manuscript. All authors contributed critically to the drafts and gave final approval for publication.

