## Supplemental files for "Root trait responses to drought depend on plant functional group"

#### Supporting information

**Table S1.** Multivariate analysis of variance of leaf and root traits in response to drought. Water availability (i.e., drought), plant species, and their respective interactions were considered as fixed factors. F-values and significance (*, $p < 0.05$; **, $p < 0.01$; ***, $p < 0.001$; all significant values are shown in bold).

| Source of variation | MANOVA |
|---|---|
| Plant species (Ps) | **8.65*** ** |
| Water availability (Wa) | **15.28*** ** |
| Ps x Wa | **1.45** ** |

**Table S2.** Leaf, root trait and soil properties values per species growing under drought and non-drought (control) conditions. Significance was established when the non-drought and drought treatment differs ($p < 0.05$, ± 1 SE, n = 5). Specific leaf area (SLA), leaf dry matter content (LDMC), RAD (root diameter), RTD (root tissue density), SRL (specific root length), SRSA (specific root surface area).

Leaf traits

| | Shoot mass (g) | | SLA ($cm^2$ $g^{-1}$) | | LDMC (mg $g^{-1}$) | | Leaf C (g/kg) | | Leaf N (g/kg) | |
|---|---|---|---|---|---|---|---|---|---|---|
| Plant species | non-drought | drought | non-drought | drought | non-drought | drought | non-drought | drought | non-drought | drought |
| *Anthoxanthum* | 4.5 ± 0.0 | 4.3 ± 0.3 | 305.5 ± 32.8* | 195.7 ± 7.6 | 234.5 ± 7.1* | 297.7 ± 12.9 | 407.2 ± 6.8 | 375.7 ± 25.7 | 9.5 ± 0.9* | 4.7 ± 0.3 |
| *Arrhenatherum* | 3.7 ± 0.4* | 3 ± 0.4 | 324.8 ± 32.4 | 306.2 ± 12.3 | 253.8 ± 16.4* | 278.4 ± 7.6 | 420.2 ± 8.1 | 388.2 ± 24.3 | 11 ± 1.5* | 8.2 ± 1.2 |
| *Dactylis* | 5.5 ± 0.8* | 4.5 ± 0.6 | 246.9 ± 21.9 | 182.1 ± 42.4 | 236.3 ± 14.2 | 409.1 ± 131.4 | 457.8 ± 28.8* | 400.6 ± 19.9 | 8.3 ± 0.8* | 6.9 ± 0.5 |
| *F. brevipila* | 5.1 ± 0.3* | 3.2 ± 0.5 | 188.5 ± 36.1* | 139.4 ± 20.3 | 286 ± 14.3* | 348.5 ± 16.9 | 445.5 ± 4.9 | 445.4 ± 3.4 | 11.6 ± 2 | 13.9 ± 3.5 |
| *F. rubra* | 5.6 ± 0.9* | 4.1 ± 0.3 | 103.1 ± 31.8 | 113.4 ± 19.9 | 374.8 ± 91.4 | 361.2 ± 19.5 | 404.1 ± 5.3 | 399.2 ± 19.2 | 9.1 ± 1.3* | 6.1 ± 0.5 |
| *Holcus* | 6.1 ± 0.2* | 4.7 ± 0.4 | 277.3 ± 21.9 | 282.2 ± 35 | 211.1 ± 19.2 | 270.6 ± 46.4 | 400.9 ± 14.9* | 426.3 ± 6.4 | 6.9 ± 0.5 | 6.8 ± 0.4 |
| *Lolium* | 4.6 ± 0.3 | 4.9 ± 0.7 | 219.5 ± 17.5* | 163.5 ± 12.8 | 170 ± 8.3* | 223.2 ± 16.4 | 393.7 ± 3.9* | 417.1 ± 4.4 | 8.2 ± 0.7* | 10.7 ± 0.9 |
| *Poa* | 4.9 ± 0.7* | 2.9 ± 0.4 | 155.6 ± 15.9 | 140.5 ± 28.1 | 320.5 ± 20.3 | 354.2 ± 24.3 | 417.1 ± 27.6 | 409.5 ± 15.3 | 12 ± 1.2 | 12.8 ± 2.3 |
| *Achillea* | 2.4 ± 0.1 | 2.2 ± 0.2 | 137.8 ± 4.2* | 114.6 ± 9.4 | 231.7 ± 49 | 371.6 ± 93.4 | 348.1 ± 18.6* | 422.3 ± 19.8 | 14.4 ± 1.6 | 15.4 ± 1.7 |
| *Armeria* | 2.2 ± 0.6* | 1.3 ± 0.4 | 211.5 ± 23.3* | 174.1 ± 19.5 | 127.7 ± 4.4* | 163.7 ± 15.2 | 451.2 ± 14.8* | 476.9 ± 12.1 | 36.1 ± 2.2* | 33.1 ± 0.7 |
| *Artemisia* | 2.5 ± 0.3* | 1.4 ± 0.1 | 149 ± 3.9 | 151.5 ± 7.5 | 147.8 ± 10.8 | 146.1 ± 14.3 | 360.8 ± 27.7* | 383.8 ± 12.1 | 20.4 ± 1.4 | 18.4 ± 1.7 |
| *Berteroa* | 3.6 ± 0.6* | 2.7 ± 0.2 | 253.4 ± 13.7 | 235.7 ± 30.1 | 156.7 ± 6.6 | 169.2 ± 15.9 | 326.1 ± 36.3 | 347.3 ± 17.7 | 21.1 ± 3.4 | 21.7 ± 2.4 |
| *Daucus* | 3.8 ± 0.6* | 1.7 ± 0.3 | 206.8 ± 6.86* | 177.5 ± 9.5 | 176.6 ± 6.5* | 198.7 ± 6.9 | 386.2 ± 20.7 | 375.6 ± 18.3 | 12.7 ± 1.2* | 10.6 ± 0.8 |
| *Galium* | 4.0 ± 0.9* | 2.7 ± 0.3 | 194.9 ± 10.8* | 139.4 ± 10.1 | 189.3 ± 10.7* | 220.6 ± 17.6 | 384.3 ± 17.4 | 396.2 ± 12.4 | 15.9 ± 1.1* | 11.5 ± 1 |
| *Hieracium* | 3.9 ± 0.5* | 3 ± 0.2 | 237 ± 30.7* | 160.1 ± 19.4 | 98.7 ± 22.6* | 163.9 ± 16.4 | 471.7 ± 25.2* | 406.1 ± 21.7 | 23.2 ± 2.1* | 13 ± 1.1 |
| *Hypericum* | 2.7 ± 0.2* | 1.7 ± 0.2 | 471.8 ± 90.9* | 276 ± 20.2 | 138.6 ± 19.7* | 195.4 ± 2.5 | 454.8 ± 20 | 443.2 ± 14.9 | 26.1 ± 2.7* | 17.1 ± 0.6 |
| *Plantago* | 5.1 ± 1.2* | 3.4 ± 0.4 | 185.9 ± 17.9 | 168.8 ± 33.7 | 165.6 ± 11.1 | 187.3 ± 13.5 | 368.1 ± 14.4 | 348.9 ± 31.3 | 9.9 ± 1.4 | 10.4 ± 1.6 |
| *Potentilla* | 3.8 ± 0.8* | 3.3 ± 0.8 | 110.5 ± 11.1* | 78.8 ± 2.4 | 351.5 ± 24.5* | 406.1 ± 10.3 | 465.6 ± 30.3 | 459.9 ± 6.8 | 17.7 ± 3* | 14.5 ± 0.9 |
| *Ranunculus* | 2.2 ± 0.2 | 1.7 ± 0.3 | 196.2 ± 19.1* | 167.9 ± 8.6 | 189.4 ± 9.3 | 200.9 ± 9.1 | 405 ± 11.6 | 404.8 ± 6.7 | 17 ± 3.2* | 14 ± 0.5 |
| *Rumex* | 3.4 ± 0.6* | 1.8 ± 0.4 | 178.7 ± 11.4 | 229.4 ± 34.6 | 169.6 ± 7.5* | 139.6 ± 8.2 | 390.1 ± 15.7 | 389.3 ± 7.3 | 12.6 ± 0.9 | 14.3 ± 2 |
| *Silene* | 3.9 ± 0.5* | 2.7 ± 0.4 | 232 ± 45.9* | 175.3 ± 10.7 | 121.5 ± 11.7 | 154.3 ± 13 | 384.4 ± 15.5* | 405.7 ± 5.3 | 20.3 ± 1.4 | 16.4 ± 1.8 |
| *Medicago* | 9.3 ± 1.3* | 5.4 ± 0.8 | 278.1 ± 48.4* | 223.6 ± 19.1 | 174.9 ± 57.5* | 208.9 ± 14.6 | 431.4 ± 12 | 449.6 ± 14.1 | 38.4 ± 1.7 | 37.7 ± 1.1 |
| *Trifolium* | 16.5 ± 2.3* | 7.6 ± 0.6 | | | | | 411.5 ± 48.2 | 455 ± 19.7 | 38.4 ± 5.6 | 35.7 ± 1.5 |
| *Vicia* | 4.3 ± 0.9* | 2.5 ± 0.3 | 404.3 ± 30.1* | 350.6 ± 24.1 | 198.2 ± 23.3* | 223.6 ± 11 | 421.5 ± 16.6 | 394.7 ± 22.2 | 36.3 ± 2.6* | 31 ± 2.4 |

### Root traits

| | Root biomass (g) | | RAD (mm) | | RTD (mg $cm^{-3}$) | | SRL (cm $mg^{-1}$) | | SRSA (cm2 $mg^{-1}$) | | Root C (g/Kg) | | Root N (g/Kg) | |
|---|---|---|---|---|---|---|---|---|---|---|---|---|---|---|
| Plant species | non-drought | Drought | non-drought | Drought | non-drought | Drought | non-drought | Drought | non-drought | Drought | non-drought | Drought | non-drought | Drought |
| *Anthoxanthum* | 4.1 ± 0.9 | 3.4 ± 0.8 | 0.12± 0.01* | 0.2 ± 0.02 | 267 ± 10.5* | 188 ± 30.2 | 33.5 ± 2.1* | 28.1 ± 1.7 | 1.3 ± 0* | 1.4 ± 0.1 | 395.5 ± 56.7 | 371.7 ± 26.3 | 5.5 ± 1.1 | 9.4 ± 3.2 |
| *Arrhenatherum* | 4.5 ± 0.8* | 7.9 ± 0.5 | 0.25± 0.01* | 0.3 ± 0.02 | 335.9 ± 37.9* | 172.5 ± 8.9 | 6.8 ± 1.1 | 8.2 ± 0.8 | 0.5 ± 0.1* | 0.8 ± 0 | 432.7 ± 90.1 | 411.8 ± 76.3 | 5.9 ± 1.2 | 5.8 ± 1.2 |
| *Dactylis* | 2.7 ± 0.8* | 4.8 ± 0.6 | 0.17± 0.01* | 0.2 ± 0.01 | 296.5 ± 24.4* | 210.4 ± 17 | 16.2 ± 1.8* | 12.9 ± 1.4 | 0.8 ± 0.1 | 0.9 ± 0.1 | 367.5 ± 9.8 | 400.1 ± 28.2 | 4.6 ± 0.4 | 4.9 ± 0.5 |
| *F. brevipila* | 3.2 ± 0.3* | 1.7 ± 0.6 | 0.25± 0.03* | 0.2 ± 0.01 | 245.5 ± 78.5* | 180.6 ± 7.7 | 10.2 ± 2.2* | 15.2 ± 1.1 | 0.8 ± 0.2* | 1 ± 0.1 | 409 ± 18.1 | 390.9 ± 18.4 | 6.8 ± 0.3 | 8.4 ± 1 |
| *F. rubra* | 5.2 ± 0.8* | 4.5 ± 0.4 | 0.21± 0.03 | 0.2 ± 0.01 | 189.2 ± 20.3* | 226.5 ± 14.3 | 17.2 ± 2.9* | 14.8 ± 1 | 1.1 ± 0.1* | 0.9 ± 0 | 396 ± 22.3 | 390.1 ± 9.7 | 5.6 ± 0.5* | 5.2 ± 0.2 |
| *Holcus* | 5.6 ± 1.2 | 5.4 ± 0.6 | 0.25± 0.01 | 0.3 ± 0.01 | 158.4 ± 26.3* | 123.4 ± 7.6 | 13.9 ± 1.1 | 16.1 ± 1.1 | 1.1 ± 0.1* | 1.3 ± 0.1 | 376.4 ± 15 | 362.2 ± 29.6 | 4.1 ± 0.3 | 4.6 ± 0.4 |
| *Lolium* | 6.9 ± 1* | 5.4 ± 0.8 | 0.2± 0.01* | 0.2 ± 0.01 | 180.5 ± 5.5* | 159.1 ± 9.8 | 17.2 ± 1.2 | 13.8 ± 2.1 | 1.1 ± 0 | 1 ± 0.1 | 299.8 ± 20.7 | 305.7 ± 18.8 | 4.7 ± 0.4 | 4.9 ± 0.4 |
| *Poa* | 3.9 ± 0.5* | 2.4 ± 0.5 | 0.17± 0.01* | 0.2 ± 0 | 150.9 ± 15.4 | 156 ± 7.5 | 30 ± 4.5* | 25.1 ± 0.7 | 1.6 ± 0.2* | 1.4 ± 0 | 403.5 ± 10.5* | 366.3 ± 20.9 | 5.8 ± 0.3 | 6.2 ± 0.5 |
| *Achillea* | 1.1 ± 0.5* | 2.7 ± 0.4 | 0.25± 0.02* | 0.3 ± 0.02 | 436.7 ± 31.7* | 310.5 ± 26.9 | 5.2 ± 0.9 | 5.1 ± 0.3 | 0.4 ± 0.1* | 0.5 ± 0 | 417.3 ± 7.7 | 425.9 ± 11.4 | 10.6 ± 0.9* | 7 ± 0.6 |
| *Armeria* | 0.2 ± 0.1 | 0.2 ± 0 | 0.48± 0.16 | 0.5 ± 0.12 | 261.2 ± 60.2 | 393 ± 59.1 | 5.8 ± 2.6* | 2.1 ± 0.6 | 0.5 ± 0.1* | 0.3 ± 0 | 407.1 ± 9.9 | 373.2 ± 28.5 | 18.7 ± 1 | 18.7 ± 3 |
| *Artemisia* | 1.5 ± 0.1 | 1.8 ± 0.5 | 0.42± 0.08 | 0.4 ± 0.06 | 366.5 ± 65.8 | 370.1 ± 30.6 | 4.6 ± 2.4* | 2.8 ± 0.7 | 0.4 ± 0.1* | 0.3 ± 0 | 384 ± 15.6* | 416.2 ± 5 | 8.9 ± 1.4 | 9.3 ± 2 |
| *Berteroa* | 0.8 ± 0.1 | 0.7 ± 0.1 | 0.3± 0.06* | 0.2 ± 0.01 | 378.8 ± 28 | 373.3 ± 42.5 | 6.2 ± 2.7 | 7.3 ± 1.2 | 0.4 ± 0.1 | 0.5 ± 0.1 | 385.2 ± 7.8 | 392.4 ± 40.2 | 16.3 ± 1.1 | 12.7 ± 3.4 |
| *Daucus* | 2.9 ± 0.4* | 3.4 ± 0.1 | 0.17± 0.01* | 0.2 ± 0.01 | 390.9 ± 91.1* | 290.9 ± 13 | 15.7 ± 3.8* | 11.9 ± 1.2 | 0.8 ± 0.2 | 0.7 ± 0 | 434.6 ± 14.5 | 461.7 ± 25.6 | 5.4 ± 0.2* | 6.2 ± 0.5 |
| *Galium* | 3.5 ± 2.2* | 2 ± 0.4 | 0.26± 0.03 | 0.3 ± 0.04 | 308.5 ± 25.6 | 364 ± 40.7 | 7 ± 1.2 | 5 ± 1.5 | 0.5 ± 0.1* | 0.4 ± 0.1 | 329.3 ± 15.7* | 418.4 ± 11.6 | 7.2 ± 0.6 | 7.2 ± 0.4 |
| *Hieracium* | 0.6 ± 0.2 | 0.9 ± 0.2 | 0.28± 0.01 | 0.3 ± 0.01 | 312.1 ± 19.9* | 209.7 ± 6.4 | 5.6 ± 0.7* | 8.1 ± 0.7 | 0.5 ± 0 | 0.7 ± 0 | 333.6 ± 23.6* | 384.4 ± 7.9 | 12 ± 0.8* | 7.4 ± 0.5 |
| *Hypericum* | 1.5 ± 0.4* | 2.5 ± 0.3 | 0.31± 0.02 | 0.3 ± 0.02 | 298.3 ± 27.3 | 324.8 ± 38.5 | 5.1 ± 0.8 | 4.4 ± 0.9 | 0.5 ± 0.1 | 0.4 ± 0.1 | 370 ± 13.2* | 432.2 ± 6.4 | 7.2 ± 1.1* | 7.9 ± 0.1 |
| *Plantago* | 3.4 ± 0.7 | 4.6 ± 0.7 | 0.26± 0.01* | 0.3 ± 0.02 | 233.6 ± 60.6* | 155.1 ± 14 | 8.8 ± 1.8 | 9.2 ± 1.2 | 0.7 ± 0.1 | 0.9 ± 0.1 | 342.3 ± 13.7* | 367.7 ± 13.3 | 6.8 ± 0.6 | 7.2 ± 1.4 |
| *Potentilla* | 0.9 ± 0.2 | 1.1 ± 0.3 | 0.27± 0.03 | 0.3 ± 0.03 | 668.4 ± 86.3* | 517 ± 28.7 | 3.9 ± 1.7 | 3.3 ± 0.6 | 0.3 ± 0.1 | 0.3 ± 0 | 437.2 ± 28.4 | 444.6 ± 44.1 | 10.6 ± 0.7 | 11 ± 1.1 |
| *Ranunculus* | 4.3 ± 0.9* | 3.3 ± 0.2 | 0.31± 0.03 | 0.3 ± 0.04 | 238 ± 30 | 287.6 ± 53.4 | 6.7 ± 1.5 | 4.8 ± 1.2 | 0.6 ± 0.1 | 0.5 ± 0.1 | 393.9 ± 4.4 | 384.1 ± 23.5 | 8 ± 3.7* | 6.2 ± 0.4 |
| *Rumex* | 7 ± 1.3* | 4.1 ± 0.5 | 0.41± 0.13* | 0.9 ± 0.12 | 2168.2±279.1* | 1366.4 ± 156.2 | 0.7 ± 0.2* | 0.2 ± 0 | 0.1 ± 0* | 0 ± 0 | 310.2 ± 78.5* | 475.8 ± 3.3 | 6.5 ± 0.2* | 9 ± 1 |
| *Silene* | 4.4 ± 0.5 | 5.4 ± 0.7 | 0.48± 0.04* | 0.4 ± 0.04 | 555.3 ± 34.1 | 628.2 ± 43.4 | 1.1 ± 0.2 | 1.8 ± 0.4 | 0.2 ± 0 | 0.2 ± 0 | 404.6 ± 8.4 | 388.4 ± 17 | 5.5 ± 0.6* | 4.6 ± 0.1 |
| *Medicago* | 1 ± 0.1 | 0.8 ± 0.2 | 0.24± 0.01* | 0.3 ± 0.01 | 359.9 ± 21.3 | 347.2 ± 45.5 | 6.3 ± 0.6 | 5.4 ± 1 | 0.5 ± 0 | 0.5 ± 0.1 | 374.3 ± 11.1 | 353.2 ± 15.1 | 17.6 ± 1.2 | 19.3 ± 1.6 |
| *Trifolium* | 1.4 ± 0.2* | 0.9 ± 0.2 | 0.31± 0.02* | 0.2 ± 0.01 | 353.2 ± 21.6 | 366.3 ± 35.3 | 4 ± 0.6* | 7.2 ± 1.4 | 0.4 ± 0* | 0.5 ± 0.1 | 362.7 ± 9.5 | 398.7 ± 20.7 | 17.4 ± 0.5* | 20.4 ± 0.4 |
| *Vicia* | 1.8 ± 0.5* | 1.1 ± 0.3 | 0.52± 0.08 | 0.5 ± 0.03 | 323.4 ± 21.1* | 418.3 ± 13.9 | 2.2 ± 1* | 1.3 ± 0.2 | 0.3 ± 0.1* | 0.2 ± 0 | 407.1 ± 9.4 | 422.8 ± 11.7 | 39 ± 2.9* | 32.8 ± 3.3 |

### Soil properties

| | Soil C (g/Kg) | | Soil N (g/Kg) | |
|---|---|---|---|---|
| Plant species | non-drought | Drought | non-drought | Drought |
| *Anthoxanthum* | 7.34 ± 0.18 | 7.67 ± 0.17 | 0.66 ± 0.02 | 0.68 ± 0.01 |
| *Arrhenatherum* | 7.06 ± 0.20 | 9.00 ± 1.49 | 0.62 ± 0.03 | 0.77 ± 0.13 |
| *Dactylis* | 9.46 ± 1.68 | 7.15 ± 0.31 | 0.77 ± 0.17 | 0.70 ± 0.03 |
| *F. brevipila* | 7.66 ± 0.42 | 7.40 ± 0.27 | 0.62 ± 0.03 | 0.63 ± 0.02 |
| *F. rubra* | 7.43 ± 0.14 | 7.15 ± 0.16 | 0.66 ± 0.03 | 0.62 ± 0.02 |
| *Holcus* | 7.20 ± 0.11 | 6.74 ± 0.29 | 0.67 ± 0.001 | 0.62 ± 0.02 |
| *Lolium* | 6.48 ± 0.65 | 8.03 ± 0.33 | 0.61 ± 0.06 | 0.70 ± 0.02 |
| *Poa* | 6.54 ± 0.19 | 6.98 ± 0.23 | 0.68 ± 0.04 | 0.71 ± 0.05 |
| *Achillea* | 7.58 ± 0.32 | 7.83 ± 0.25 | 0.65 ± 0.02 | 0.68 ± 0.02 |
| *Armeria* | 7.16 ± 0.17 | 7.12 ± 0.25 | 0.71 ± 0.01 | 0.71 ± 0.02 |
| *Artemisia* | 7.83 ± 0.35 | 7.22 ± 0.27 | 0.63 ± 0.03 | 0.63 ± 0.01 |
| *Berteroa* | 7.92 ± 0.37* | 11.84 ± 2.09 | 0.66 ± 0.03* | 1.06 ± 0.20 |
| *Daucus* | 9.87 ± 2.42 | 8.53 ± 1.43 | 0.80 ± 0.16 | 0.69 ± 0.13 |
| *Galium* | 7.56 ± 0.32 | 6.87 ± 0.40 | 0.66 ± 0.03 | 0.67 ± 0.05 |
| *Hieracium* | 7.14 ± 0.21 | 7.51 ± 0.22 | 0.63 ± 0.01 | 0.65 ± 0.03 |
| *Hypericum* | 8.23 ± 0.43 | 6.94 ± 0.33 | 0.69 ± 0.02 | 0.64 ± 0.01 |
| *Plantago* | 7.13 ± 0.06 | 7.98 ± 0.86 | 0.64 ± 0.01 | 0.65 ± 0.01 |
| *Potentilla* | 7.75 ± 0.25 | 7.26 ± 0.24 | 0.66 ± 0.02 | 0.65 ± 0.01 |
| *Ranunculus* | 7.32 ± 0.15 | 6.95 ± 0.15 | 0.71 ± 0.04 | 0.65 ± 0.01 |
| *Rumex* | 7.05 ± 0.18 | 7.53 ± 2.41 | 0.63 ± 0.01 | 0.69 ± 0.19 |
| *Silene* | 5.97 ± 0.96 | 9.14 ± 1.41 | 0.61 ± 0.08* | 0.94 ± 0.17 |
| *Medicago* | 7.52 ± 0.16 | 7.54 ± 0.10 | 0.76 ± 0.06 | 0.72 ± 0.01 |
| *Trifolium* | 10.37 ± 2.90 | 8.29 ± 1.16 | 0.96 ± 0.29 | 0.87 ± 0.11 |
| *Vicia* | 8.57 ± 1.31 | 7.41 ± 0.19 | 0.79 ± 0.13 | 0.65 ± 0.02 |

**Table S3.** Leaf and root trait values per plant functional group growing under non-drought (control) conditions. Significance was established at $p < 0.05$, means ± 1 SE, n = 5). Specific leaf area (SLA), leaf dry matter content (LDMC), RAD (root diameter), RTD (root tissue density), SRL (specific root length), SRSA (specific root surface area).

| | Grasses | Herbs | Legumes |
|---|---|---|---|
| Leaf traits | | | |
| Shoot mass (g) | 5.0 ± 0.26 b | 3.3 ± 0.24 c | 9.9 ± 3.39 a |
| SLA ($cm^2$ $g^{-1}$) | 227.7 ± 26.94 b | 212.7 ± 24.33 b | 341.2 ± 63.1 a |
| LDMC (mg $g^{-1}$) | 260.9 ± 22.85 a | 174.2 ± 17.58 b | 186.6 ± 11.65 b |
| Leaf C (g/kg) | 418.3 ± 7.94 a | 399.7 ± 13.05 a | 421.5 ± 5.74 a |
| Leaf N (g/kg) | 9.6 ± 0.64 c | 19.0 ± 1.91 b | 37.7 ± 0.7 a |
| Root traits | | | |
| Root biomass (g) | 4.5 ± 0.48 a | 2.5 ± 0.55 b | 1.4 ± 0.23 b |
| RAD (mm) | 0.2 ± 0.02 b | 0.3 ± 0.03 a | 0.4 ± 0.08 a |
| RTD (mg $cm^{-3}$) | 228 ± 24.17 b | 509 ± 142.57 a | 345.5 ± 11.22 a |
| SRL (cm $mg^{-1}$) | 18.1 ± 3.25 a | 5.9 ± 1.02 b | 4.2 ± 1.19 b |
| SRSA ($cm^2$ $mg^{-1}$) | 1.04 ± 0.12 a | 0.5 ± 0.05 b | 0.4 ± 0.06 b |
| Root C (g/Kg) | 385.1 ± 14.05 a | 380.7 ± 11.42 a | 381.4 ± 13.29 a |
| Root N (g/Kg) | 5.4 ± 0.30 c | 9.5 ± 1.14 b | 24.7 ± 7.17 a |

**Table S4.** Pearson correlation between shoot mass and root traits of plants growing under drought and non-drought conditions. Positive values indicate a positive correlation. Significance established at $p<0.05$ in bold. n=118. RDA and root mass correspond to their sqrt value while SRL and RTD to their log value. Comparison between drought and non-drought treatment was done through a linear model analyses using the absolute Pearson values.

| | Non-drought | | Drought | |
|---|---|---|---|---|
| | Pearson | p-value | Pearson | p-value |
| Root mass | 0.26 | 0.0042 | 0.21 | 0.0247 |
| RDA | -0.22 | 0.0192 | -0.35 | 0.0001 |
| RTD | -0.08 | 0.3982 | -0.31 | 0.0008 |
| SRL | 0.16 | 0.0775 | 0.38 | <0.0001 |
| SRSA | 0.16 | 0.0893 | 0.36 | 0.0001 |
| Root C | -0.12 | 0.1939 | -0.30 | 0.0010 |
| Root N | 0.04 | 0.7037 | 0.02 | 0.8188 |

Comparison between drought and non-drought (Pearson values)

| Factor | F-value | p-value |
|---|---|---|
| Water treatment | 5.22 | 0.041 |

| Water treatment | Mean | E.E. | | |
|---|---|---|---|---|
| drought | 0.28 | 0.04 | A | |
| control | 0.15 | 0.04 | | B |

**Table S5.** Magnitude of the drought effect on leaf and root traits for each plant functional group. RII compares the magnitude of the effect of drought vs non-drought treatment. Negative values indicate higher trait response in non-drought than in drought treatment and positive values indicate the opposite. RII values different significantly from 0 (*,**, *** p < 0.1, 0.05, 0.01).

| RII | All species | Grasses | Herbs | Legumes |
|---|---|---|---|---|
| **Soil properties** | | | | |
| Soil C | -0.01 | 0.01 | 0.01 | **-0.07*** ** |
| Soil N | -1.6E-03 | 0.08 | -0.01 | **-0.07*** ** |

**Figure S1.** Mean and standard deviation of traits for the whole set of plant species. The values are scaled for each plant trait. See that the maximum value corresponds to that used in the figure 5.

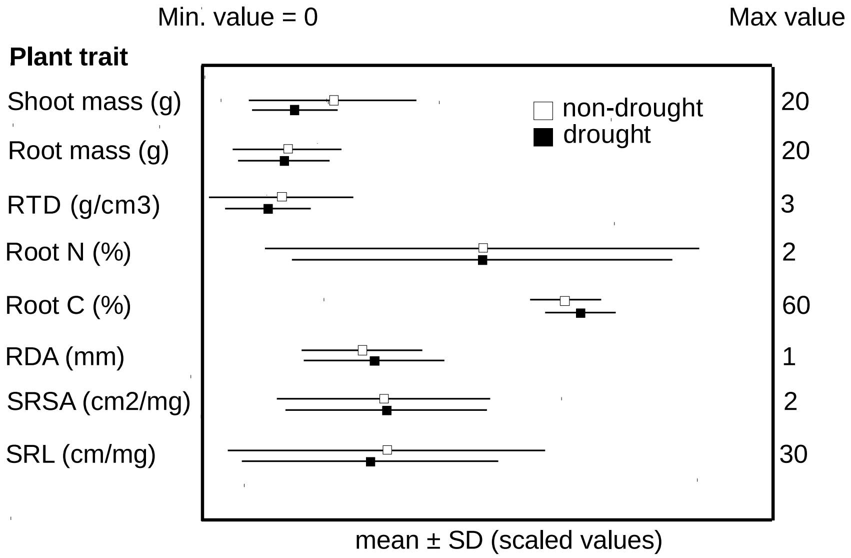

**Figure S2.** Relationships between different root traits and shoot mass under drought and non-drought treatments. a) root diameter (RDA), b) root tissue density (RTD), c) specific root length SRL) and specific root surface area (SRSA). Solid lines indicate regression lines, and shaded areas indicate 95% confidence intervals. **P* < 0.05, ***P* < 0.01, and ****P* < 0.001.

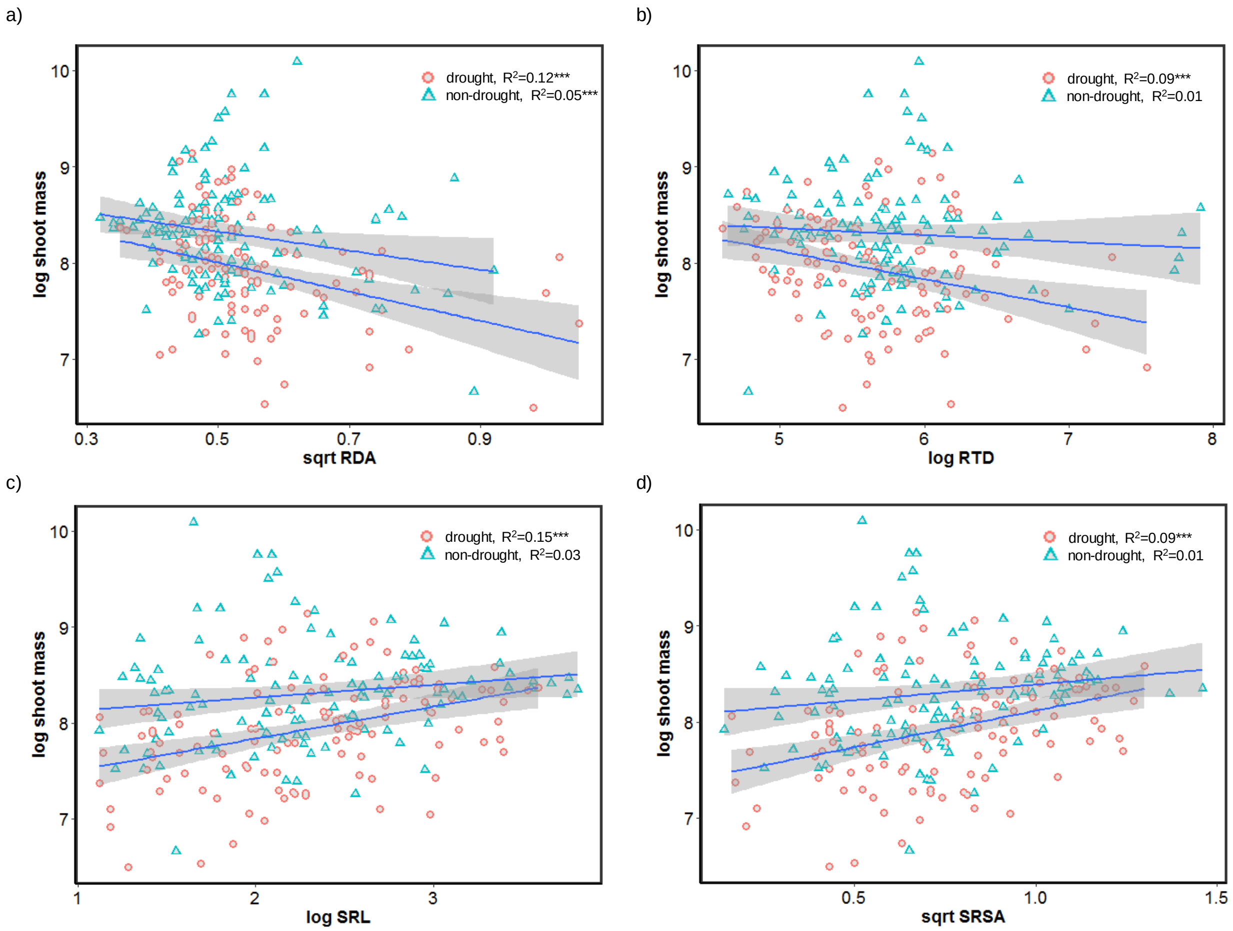

**Figure S3.** Principal component analysis of leaf (a) and root (b) traits associated to 24 plant species belonging to different plant functional groups growing under drought conditions. Plant functional groups: Grasses (square symbols), herbs (circle symbols) and legumes (triangle symbols). Plant traits: SLA (specific leaf area), LDMC (leaf dry matter content), RDA (root diameter), RTD (root tissue density), SRL (specific root length), SRSA (specific root surface area). Plant species: *Anthoxanthum* (blue 2), *Berteroa* (light yellow), *Daucus* (light green), *F.brevipila* (light gray), *Achillea* (salmon), *Potentilla* (deepskyblue4), Plantago (yellow), *Hieracium* (darkseagreen1), *Artemisa* (azure 2), *Holcus* (pink), *Arrhenaterum* (purple), *Vicia* (golden), *Hypericum* (chartreuse)*, F.rubra* (azure 3), *Galium* (orange), *Poa* (dark blue), *Silene* (dark golden), *Trifolium* (green), it was not included in the graph 'a', *Dactylis* (azure 4), *Rumex* (light pink), *Medicago* (dark purple), *Ranunculus* (Cyan 4), *Armeria* (Dark grey). n=5.

a)

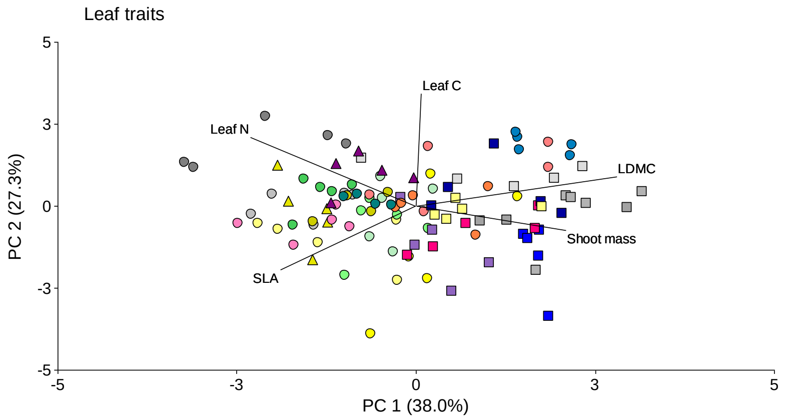


b)

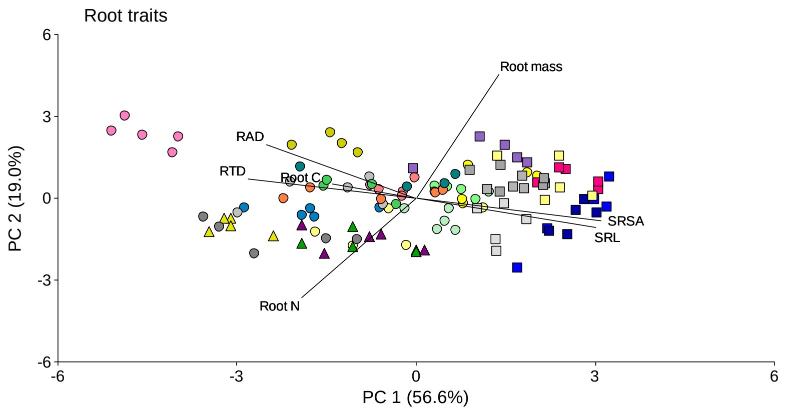

**Figure S4.** Root length of herb plants with a diameter lower than 0.1mm. Means ± EE

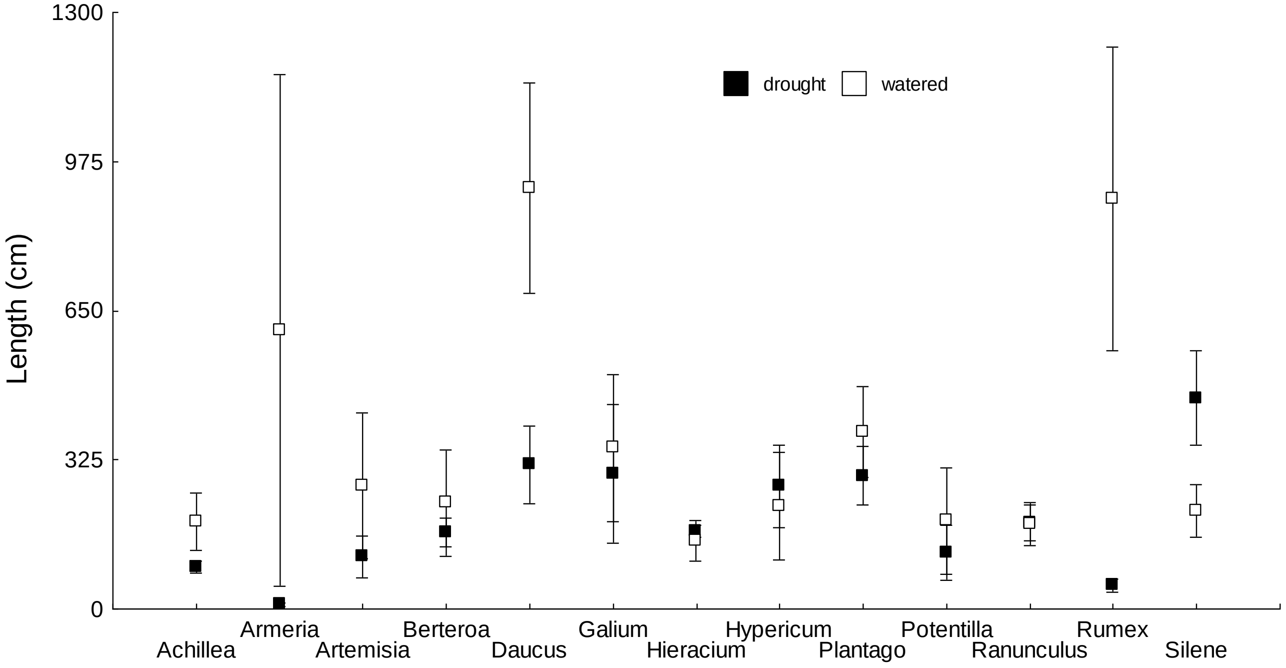
